# Postsynaptic potential energy as determinant of synaptic plasticity

**DOI:** 10.1101/2020.08.26.269290

**Authors:** Huan-Wen Chen, Li-Juan Xie, Yi-Jun Wang, Hang Zhang

## Abstract

Metabolic energy can be used as a unified principle to control neuronal activity. However, whether and how metabolic energy alone can determine the outcome of synaptic plasticity remains unclear. In this study, a computational model of synaptic plasticity that is completely determined by energy is proposed. A simple quantitative relationship between synaptic plasticity and postsynaptic potential energy is established. Synaptic weight is directly proportional to the difference between the baseline potential energy and the suprathreshold potential energy and is constrained by the maximum energy supply. Results show that the energy constraint improves the performance of synaptic plasticity and avoids setting the hard boundary of synaptic weights. With the same set of model parameters, our model can reproduce several classical experiments in homo and heterosynaptic plasticity. The proposed model can explain the interaction mechanism of Hebbian and homeostatic plasticity at the cellular level, thereby providing a new way to deeply understand the characteristics of learning and memory.

## Introduction

Although the brain accounts for only 2% of the body mass, it consumes 20% of the resting metabolic energy produced by the whole body (Attwell and Laughlin, 2001; Harris et al., 2012). Within the brain, neurons utilize 75%–80% of this energy, and the remainder is used by the neighboring glial cells. Housekeeping tasks use 25% of the total neuronal energy. Maintaining resting membrane potential (15%), firing action potentials (16%), and synaptic transmission (44%) compose the energetically most expensive processes (Harris et al., 2012; Howarth et al., 2012). Thus, the majority of energy used by neurons is locally consumed at the synapse. In addition to the energetic costs of neural computation and transmission, experimental evidence indicates that synaptic plasticity is metabolically demanding (Mery and Kawecki, 2005; Placais and Preat, 2013; Jaumann et al., 2013; Placais et al., 2017). The energy cost of synaptic plasticity is estimated based on the neurophysiological and proteomic data of rat brain depending on the level of protein phosphorylation; this cost constitutes a small fraction of the energy used for fast excitatory synaptic transmission, which is typically 4.0%–11.2% (Karbowski, 2019). However, the quantitative relationship between the changes in synaptic weights (potentiation or depression) and energy consumption remains unclear.

Considering the consistency with corresponding experiments, a large number of synaptic plasticity models have been established. These models are mainly biophysical models based on calcium hypothesis (Shouval, et al., 2002; Graupner and Brunel, 2012) and phenomenological models based on pre and postsynaptic spikes or voltage (Bienenstock, et al., 1982; Pfister and Gerstner, 2006; Clopath, et al., 2010). Although these models are successful in experimental reproduction, they ignore the role of metabolic energy. Growing evidence suggests that metabolic energy may be a unifying principle governing neuronal activities (Laughlin, 2001; Niven and Laughlin, 2008; Hasenstaub et al., 2010; Yu and Yu, 2017), thereby naturally leading people to focus on the relationship between metabolic energy and synaptic plasticity in recent years. Sacramento et al. (2015) showed that unbalanced synaptic plasticity rules can lead to sparse connectivity and energy efficient computation. Li and van Rossum (2020) assumed that the metabolic energy for every modification of a synaptic weight is proportional to the amount of change, regardless if this is positive or negative. They proposed a synaptic caching algorithm based on this assumption. The proposed algorithm can enhance energy efficiency manifold by precisely balancing labile forms of synaptic plasticity with many stable forms. However, energy is expressed by synaptic weights in these studies. Whether synaptic plasticity can be fully quantified by energy remains unclear.

Potential energy is stored in transmembrane ion gradients. When postsynaptic neurons are stimulated by external stimuli (such as synaptic input), the changes in gating state, channel conductance, and current are driven by the energy stored in the Na^+^ and K^+^ gradients, and no adenosine triphosphate (ATP) is consumed in this process. These gradients and stored potential energy are partially depleted and must be actively restored. The change in postsynaptic potential energy indirectly reflects the consumption or supply of metabolic energy because the active recovery of potential energy needs ATP. In this study, we express the postsynaptic potential energy as the integral of the product of postsynaptic membrane potential and the postsynaptic membrane current density on neural activity time. The potential energy with membrane potential lower than a certain threshold is called subthreshold potential energy. The part with membrane potential greater than the threshold is called suprathreshold potential energy. The baseline potential energy is the result of downscaling the amplitude of the subthreshold potential energy. The synaptic weights are expressed by a simple linear relationship between the subthreshold potential energy and the suprathreshold potential energy and are constrained by the energy supply. The simulation results show that the model can reproduce a series of classic synaptic plasticity experiments, indicating that our model is feasible.

## Results

### Construction of synaptic plasticity model

Our model uses postsynaptic potential energy to express the change in synaptic weights. Postsynaptic potential energy *P* is the integral of the product of postsynaptic membrane potential *V_m_* and postsynaptic membrane current density *I_m_* to neural activity time *t*, that is, 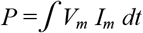 The model is constructed in four steps (**Fig. 1a**). In step (1), no neural activity occurs, and the postsynaptic neuron is in a resting state. At this time, postsynaptic potential energy *P* is the resting state potential energy *P_rest_*. Take the resting state potential energy as the reference point of potential energy, let *P* = *P_rest_* = 0. In step (2), neural activity causes potential energy *P* to deviate from the resting state potential energy. Potential energy *P* after neural activity is separated into two parts. The first part is called subthreshold potential energy *P_sub_*, and its membrane potential *V_m_* is less than threshold potential *V_th_*. The second part is called suprathreshold potential energy *P_sup_*, and its membrane potential *V_m_* is greater than *V_th_*. Thus, *P* = *P_sub_* + *P_sup_*. In step (3), the role of subthreshold potential energy in the change in synaptic weight is reduced. In particular, the subthreshold potential energy is multiplied by a constant *A_r_* between zero and one, which is called the baseline coefficient, and the baseline potential energy *P_bas_* is obtained. Thus, *P_bas_* = *A_r_ P_sub_*. In step (4), we assume that the change in synaptic weights is proportional to the difference between the baseline potential energy and the suprathreshold potential energy. Therefore, the change in synaptic weight relative to the initial synaptic weight Δ*W* is expressed as

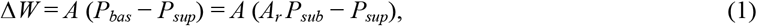

where *A* is a positive constant, which represents the linear transformation between potential energy and synaptic weight and is called the amplitude coefficient. The synaptic weights increase when the baseline potential energy is greater than the suprathreshold potential energy (**Fig. 1b**). The synaptic weights decrease when the baseline potential energy is less than the suprathreshold potential energy (**Fig. 1c**). The synaptic weights do not change when they are the same.

**Figure 1.**
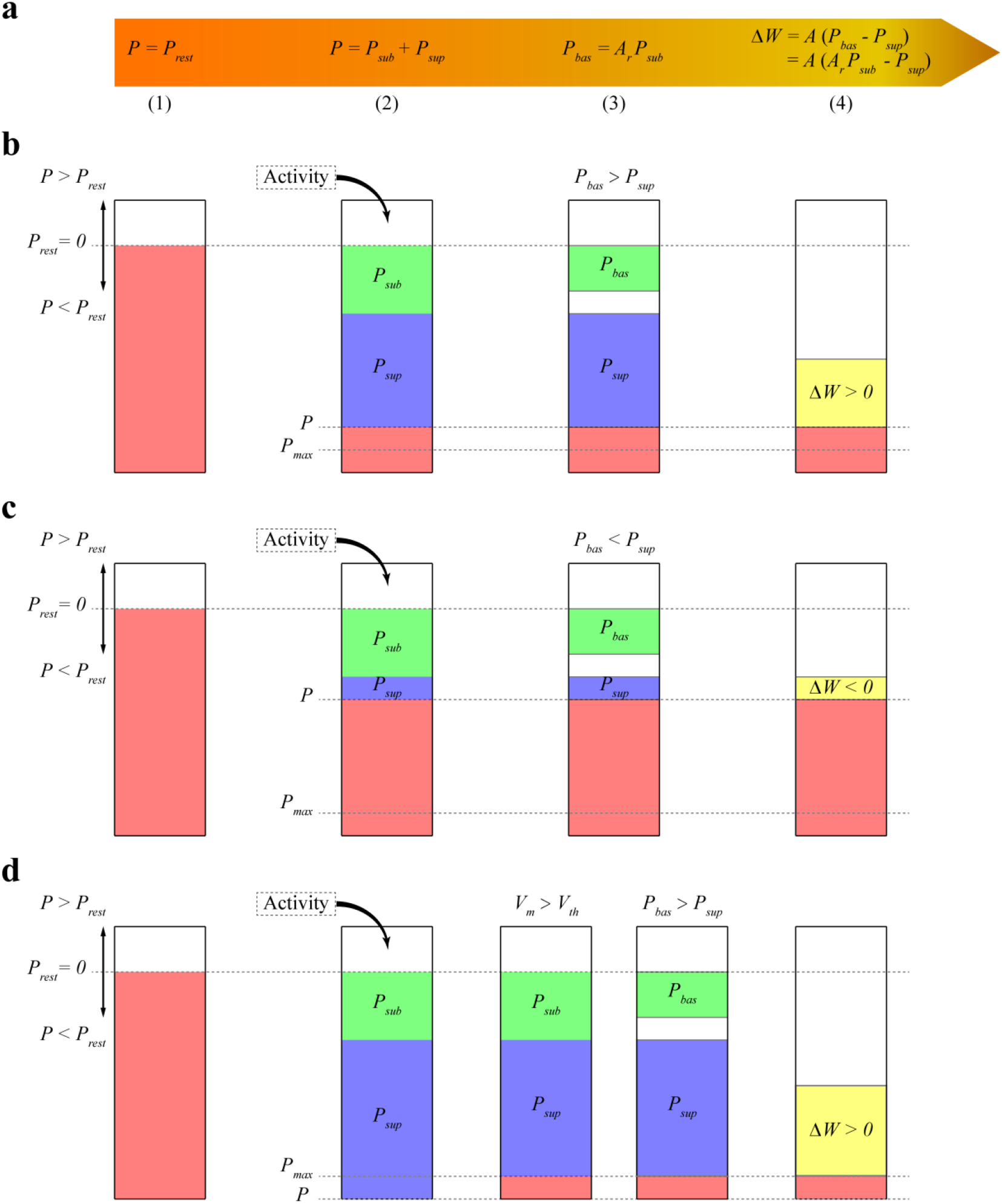
Model schematic. (**a**) Mathematical expression of synaptic plasticity model. (**b**) Calculation of synaptic weights when the baseline potential energy is greater than the suprathreshold potential energy. In resting state, the potential energy of postsynaptic membrane is assumed to be zero. The potential energy *P* of postsynaptic membrane is lower than that of resting state due to the decrease in transmembrane ion gradient. Thus, *P* < *P_rest_* is negative. Subthreshold potential energy *P_sub_* and suprathreshold potential energy *P_sup_* are negative. Baseline coefficient *A_r_* is greater than zero, so baseline potential energy *P_bas_* has the same sign as subthreshold potential energy *P_sub_* and is also negative. Here, *P_bas_* > *P_sup_*, and the synaptic weight increases. (**c**) Calculation of synaptic weights when *P_bas_* < *P_sup_*. Similar to that in (**b**), the synaptic weight decreases at this time because *P_bas_* < *P_sup_*. (**d**) Calculation of synaptic weight when the amplitude of potential energy *P* exceeds energy supply *S*. The amplitude of *P* is adjusted to the maximum potential energy *P_max_*, which is the same as *S*. Assuming that the membrane potential is greater than the threshold potential, the amplitude of *P_sup_* is reduced. The amplitude of *P_sub_* is reduced if the membrane potential is less than the threshold potential. Thus, *P_sub_* + *P_sup_* = *P* = *P_max_*.

We assume that the amplitude of postsynaptic potential energy *P* cannot be greater than the maximum energy supply *S* because the change in potential energy is constrained by energy supply. Unless otherwise specified, the energy supply in this study represents the maximum energy supply that can be provided. We call the maximum potential energy *P_max_* where its amplitude is the same as energy supply *S*, but the sign is consistent with potential energy *P*. Then *P_max_* = *S* sign (*P*), where sign is the sign function, and |*P*| ≤ *S* and |*P*| ≤ |*P_max_*|. Given that the dynamic characteristics of energy supply are unclear, we propose a simple formula for calculating the energy supply with time

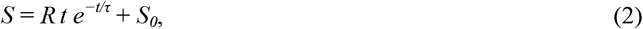

where *t* is the duration of neural activity, and *τ* is the time constant of energy supply. *R* is the energy supply per unit time, which is a constant and is called the rate of energy supply. *S_0_* is the minimum energy supply to maintain the normal function of neurons, which is a constant greater than zero. The formula represents the maximum energy that can be provided at time *t* by multiplying *Rt* of the energy supply linearly increasing with time and a damping factor *S_damp_* = *e^−t/τ^* which decreases exponentially with time. *S* = *S_0_* at *t* = 0 or *t* → ∞, the energy supply is minimum, and the energy supply is maximum at *t* = *τ*.

Potential energy *P* is adjusted to *P_max_* if its amplitude exceeds energy supply *S* (**Fig. 1d**) so that its amplitude is equal to energy supply *S*. This adjustment results in a decrease in the amplitude of suprathreshold potential energy *P_sup_* or subthreshold potential energy *P_sub_* so that their sum is equal to *P_max_*. The subthreshold potential energy is adjusted, and the suprathreshold potential energy remains unchanged when the membrane potential at time *t* is smaller than the threshold potential. The suprathreshold potential energy is adjusted, and the subthreshold potential energy remains unchanged when the membrane potential at time *t* is greater than the threshold potential. The adjustment of subthreshold potential energy causes the amplitude of baseline potential energy *P_bas_* to change in the same proportion, and the synaptic weight is controlled in a reasonable range (as shown in the next section). The specific implementations of **Equations 1** and **2** and **Fig. 1** are shown in the **Methods**.

### Determination of model parameters

In accordance with Equations (1) and (2), the model includes six parameters, namely, amplitude coefficient *A*, baseline coefficient *A_r_*, threshold potential *V_th_*, energy supply rate *R*, minimum energy supply *S_0_*, and time constant of energy supply *τ*. For the frequency-dependent pairing protocol used by Sjöström et al. (2001), we chose the final parameter by trial and error: *A* = 0.02, *A_r_* = 0.2, *V_th_* = −60 mV, *R* = 175 fJ/(μm^2^ s), *τ* = 2 s, *S_0_* = 25 fJ/μm^2^. In the next section, we introduce several classical experimental protocols of synaptic plasticity and illustrate how our model reproduces these different experimental results with the same set of parameters through potential energy and energy supply.

### Reproduction of the experimental results of homosynaptic plasticity

Nineteen distal and proximal compartments (magenta, **Fig. 2a**) were simulated in the basal dendrites of the L5 pyramidal neuron model. We followed two different experimental protocols on homosynaptic plasticity to compare our model with the experimental data. The first protocol was the classical spike timing-dependent plasticity (Markram et al., 1997; Sjöström et al., 2001; Bi and Poo, 1998). Each distal and proximal compartments were connected to one synapse. Postsynaptic spikes were induced by the injection of 1 nA and 3 ms current pulses into the soma of postsynaptic neurons. The initial synaptic weights were set to 0.5. For the study of spike frequency dependence, pairs of pre–post (**Fig. 2b**) or post–pre (**Fig. 2c**) spikes separated by 10 ms were repeated five times at different frequencies of 5 Hz up to 50 Hz with steps of 5 Hz and for 0.1 Hz. For the study of spike timing dependence (**Fig. 3a**), pairs of pre–post or post–pre spikes at 20 Hz were repeated five times for different time intervals Δ*t* (1, 2.5, 5, 7.5, 10, 12.5, 15, 17.5, and 20 ms). The computational results were multiplied by a scaling factor of 12 (60/5) to mimic 60 pairs of presynaptic and postsynaptic spikes in the experimental protocols of Sjöström et al. (2001). The study of Sjöström et al. (2001) focused on the weight change as a function of the frequency for a fixed Δ*t* in this pairing protocol. The second protocol was synaptic afferent where only presynaptic spikes were induced (Bliss and Lømo, 1973; Dudek and Bear, 1992). To test our model on a consistent set of data, we took the measurements of Dudek and Bear (1992) in the present study because sufficient quantitative information can be found in their study. Each distal and proximal compartments were connected to one synapse. The activation of the synapse connected to each compartment consisted of 20 pulses delivered by a Poisson process at input frequencies ranging from 1 Hz to 50 Hz (1, 3, 5, 10, 20, 30, 40, and 50 Hz) (**Fig. 3b**).

**Figure 2.**
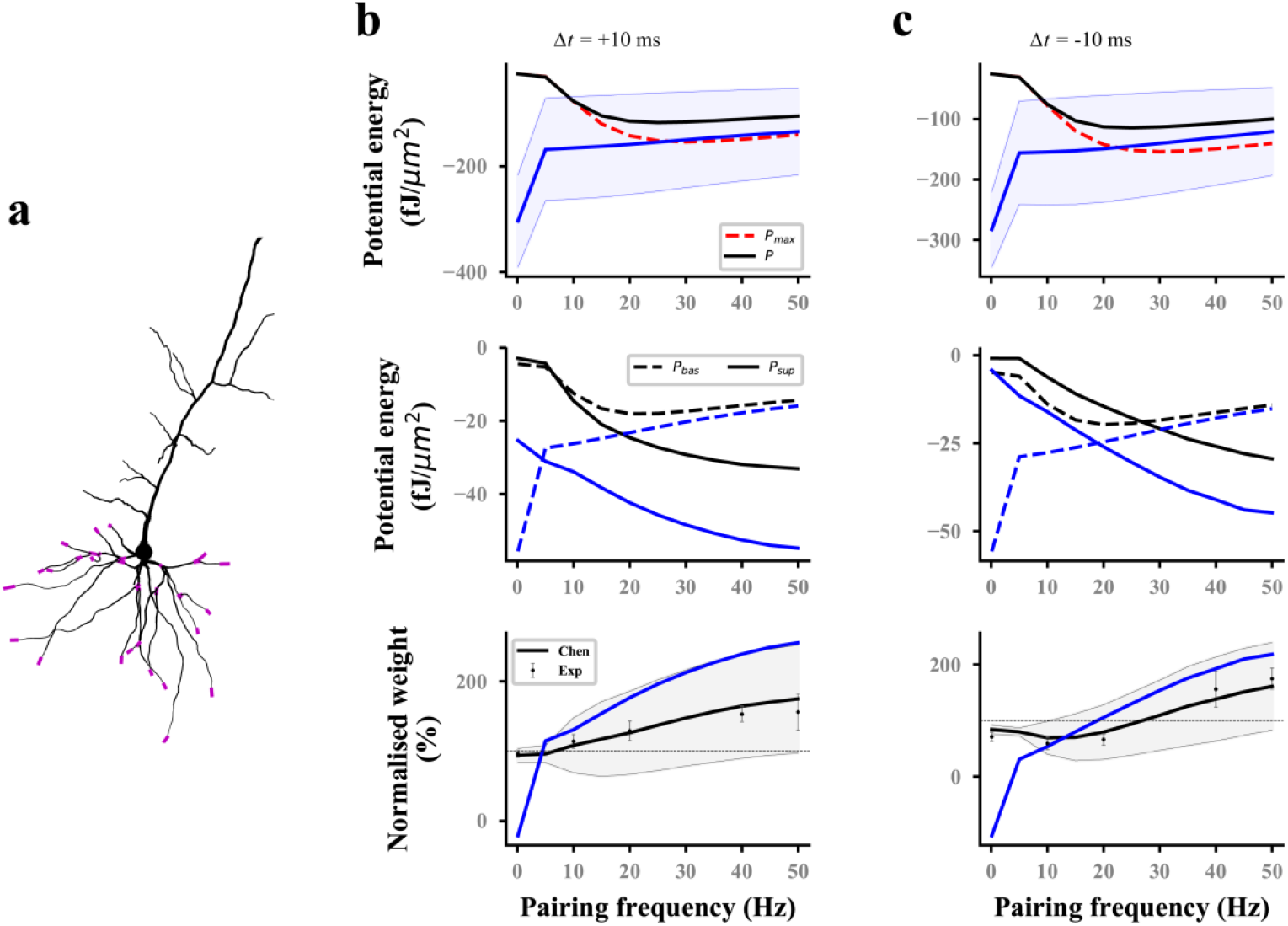
Reproducing the pairing experiment of spike frequency dependence. (**a**) Action potentials in soma are paired with either proximal or distal (magenta) synaptic activations on a thin basal branch of the L5 pyramidal neuron. (**b**, **c**) Potential energy (top), potential energy of baseline and suprathreshold (middle), and weight (bottom) change as a function of pairing repetition frequency using pairings with a time delay Δ*t* of +10 ms (pre–post, **b**) and −10 ms (post–pre, **c**). Dots with errors (bottom) represent the experimental data from Sjöström et al. (2001). The lines (solid and dashed) and the shaded regions are the mean and standard deviation (SD), respectively, over all proximal and distal compartments (magenta) shown in (**a**). The black and blue lines (solid and dashed) represent the computational results with and without energy supply constraints, respectively. For clarity, the SDs of the potential energy with constraints (black lines, top), baseline and suprathreshold energy (middle), and weights without constraints (blue lines, bottom) are not shown.

**Figure 3.**
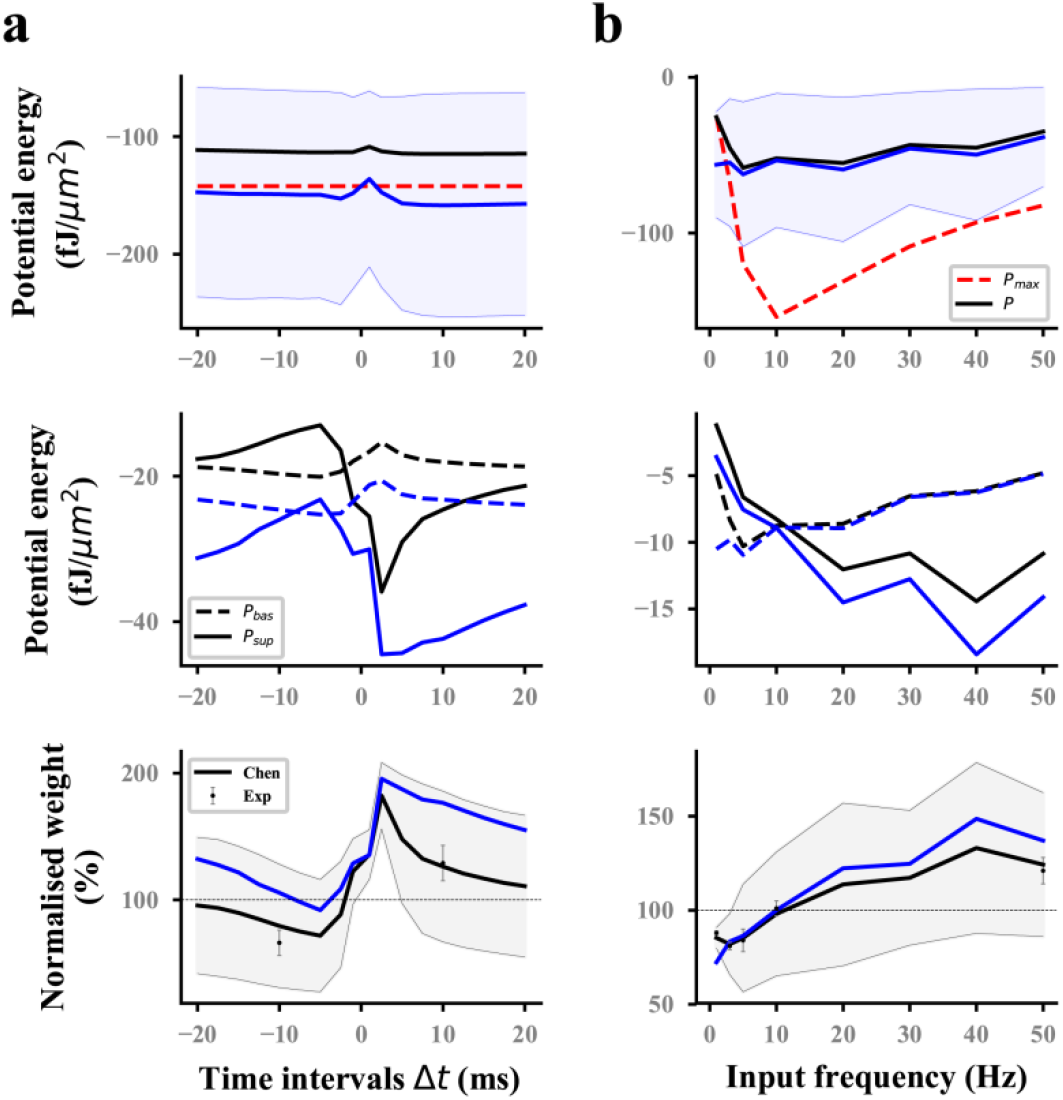
Reproducing the time-dependent pairing and synaptic afferent experiments. The lines (solid and dashed) and the shaded regions are the mean and SD, respectively, over all proximal and distal compartments (magenta) shown in **Fig. 2a**. The black and blue lines (solid and dashed) represent the computational results with and without energy supply constraints, respectively. (**a**) Potential energy (top), potential energy of baseline and suprathreshold (middle), and weight (bottom) change for different time intervals Δ*t* between pre and postsynaptic firing using 60 pre–post pairs at 20 Hz. Dots with errors (bottom) represent the experimental data from Sjöström et al. (2001). (**b**) Potential energy (top), potential energy of baseline and suprathreshold (middle), and weight (bottom) change for the synaptic afferent protocol. Dots with errors (bottom) represent the experimental data from Dudek and Bear (1992).

For the frequency-dependent pairing protocol without energy supply constraints, the amplitude of postsynaptic potential energy decreased with the increase in spike frequency (blue solid lines; **top**; **Figs. 2b,c**); this finding is consistent with the calculation results of the relationship between metabolic energy and frequency in neurons (Yi et al. 2016). With the decrease in spike frequency, the duration of neural activity increased gradually. Increasing cases were found, where the amplitude of postsynaptic potential energy without energy constraint (named as unconstrained energy) exceeded that of the maximum potential energy *P_max_* (blue shaded area under the red dotted line; **top**; **Figs. 2b,c**). The unconstrained energy with amplitude greater than that of *P_max_* was adjusted to the same as *P_max_* to obtain the postsynaptic potential energy with energy supply constraints (named as constrained energy) (black solid lines; **top**; **Figs. 2b,c**). The amplitudes of unconstrained energy in all postsynaptic compartments were greater than the amplitudes of *P_max_* when the frequency was less than 10 Hz. At this time, the amplitudes adjusted were the largest, resulting in the overlap between the constrained energy and *P_max_*. The adjustment of potential energy reduced the amplitude of baseline potential energy *P_bas_* and suprathreshold potential energy *P_sup_* (from blue dashed and solid lines to black dashed and solid lines respectively; **middle**; **Figs. 2b,c**), especially when the frequency was 0.1 Hz. The weights without energy constraints (blue solid lines; **bottom**; **Figs. 2b,c**) were adjusted to a biologically reasonable range due to the limitation of energy supply. The adjusted synaptic weights were in good agreement with the experimental data (black solid lines; **bottom**; **Figs. 2b,c**).

For the time-dependent pairing protocol (**Fig. 3a**), the neural activity time of different pairing time intervals is the same because the spike frequency was fixed at 20 Hz. Thus, the maximum potential energy does not change with the time interval (red dashed lines; **top**; **Fig. 3a**). Similar to the analysis in the previous section, the unconstrained energy with amplitude greater than *P_max_* was adjusted the same as *P_max_*. The amplitude of constrained energy (black solid lines; **top**; **Fig. 3a**) was smaller than that of the unconstrained energy (blue solid lines; **top**; **Fig. 3a**). These adjustments led to the corresponding changes in the baseline potential energy and suprathreshold potential energy (**middle**; **Fig. 3a**) and made the synaptic plasticity constrained by energy supply more consistent with the experimental results than that without energy supply constraint (**bottom**; **Fig. 3a**).

In the synaptic afferent protocol (**Fig. 3b**), increasing unconstrained energy with amplitude exceeded that of the maximum potential energy increased with the decrease in frequency when the input frequency was less than 5 Hz. The adjustment of these unconstrained energy to the maximum potential energy led to the gradual approaching and overlapping of the constrained energy and the maximum potential energy (**top**; **Fig. 3b**). If the amplitude of unconstrained energy is less than that of maximum potential energy, the constrained and unconstrained variables (i.e., potential energy, baseline and suprathreshold potential energy, and weights) should overlap because it is unnecessary to adjust the potential energy. However, the constrained and unconstrained variables did not overlap when the frequency was greater than 5 Hz although the amplitude of unconstrained energy (blue solid lines; **top**; **Fig. 3b**) was less than that of the maximum potential energy (red dashed lines; **top**; **Fig. 3b**). This phenomenon was because all the calculation results in the present study were the values at the end of neural activity. However, the adjustment of potential energy was conducted at every moment from the beginning to the end of neural activity (**Methods**). These results indicated that before the end of neural activity, several unconstrained energy were adjusted because their amplitude exceeded that of the maximum potential energy. During the neural activity, the main adjustment was reflected in the suprathreshold potential energy if the input frequency was greater than 5 Hz. However, the baseline potential energy was mainly adjusted if the input frequency was less than 5 Hz (**middle**, **Fig. 3b**). The computational results showed that our model can quantitatively reproduce the results of synaptic afferent experiment (**bottom**, **Fig. 3b**).

### Reproduction of Mexican cap-like heterosynaptic long-term depression (LTD)

Heterosynaptic plasticity can be induced at synapses that are inactive during the induction of homosynaptic plasticity (Chistiakova et al., 2015; Zenke et al., 2017). High-frequency afferent tetanization induces a Mexican hat-like profile of response amplitude changes: Homosynaptic long-term potentiation (LTP) at stimulated inputs is surrounded by heterosynaptic LTD (White et al. 1990; Royer and Paré, 2003). Each compartment on the thin basal branch of the L5 pyramidal neuron model is connected to two synapses. The initial synaptic weights were set to 0.5. To reproduce the heterosynaptic LTD, we used a similar protocol to that of Royer and Paré (2003). Homosynaptic LTP was induced with high-frequency stimuli (HFS) at the synapses connected to each distal or proximal compartment. HFS consisted of four series of 10 trains separated by 0.3 s, where each train consisting of 10 shocks (Poisson process) at 100 Hz. Each basal dendrite was divided into seven sites to compare with the experimental results. Site 0 was a compartment corresponding to homosynaptic LTP (**magenta dots**; **Fig. 4**). Other compartments connected by heterosynapses were divided into six sites in accordance with the distance from site 0 (**cyan dots**; **Fig. 4**). The value in each site (such as potential energy, weight, etc.) was the average for the values of all compartments in the site.

**Figure 4.**
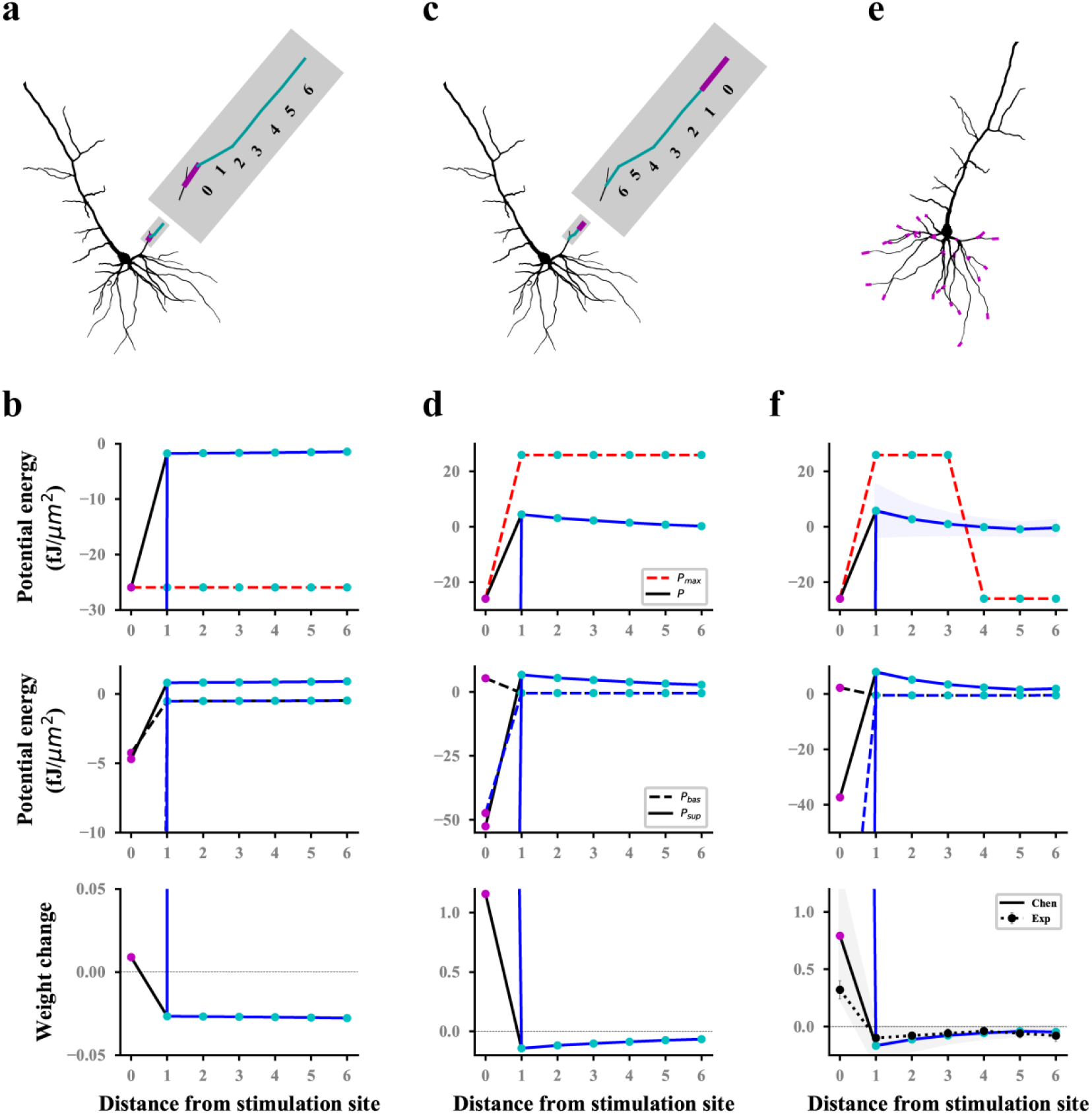
Reproducing a Mexican hat-like heterosynaptic LTD. The black and blue lines (solid and dashed) represent the computational results with and without energy supply constraints, respectively. (**a**, **b**) Schematic (**a**) of homosynaptic (magenta) and heterosynaptic (cyan) connected sites on the branch and the corresponding computational results (**b**) when stimulating the proximal synapses of a dendrite branch. (**c**, **d**) Same as (**a**) and (**b**), but for distal stimulation. (**e**, **f**) All stimulated sites (**e**) and the corresponding computational results (**f**). The lines (solid and dashed) and the shaded regions are the mean and SD, respectively, over all proximal and distal compartments (magenta) shown in (**e**). Dot line with errors (**bottom**, **f**) represent the experimental data from Royer and Paré (2003). For clarity, the SDs of the potential energy with constraints (black lines, **top**), baseline and suprathreshold energy (**middle**), and weights without constraints (blue lines, **bottom**) are not shown.

Homosynaptic LTP was induced in the synapses at the stimulation site, and heterosynaptic LTD (bottom, **Figs. 4b,d**) occurred in the nonactivated synapses of the same branch when HFS was performed at the proximal (**Fig. 4a**) or distal (**Fig. 4c**) in a thin branch. The statistical results for proximal and distal stimulation in all basal branches (**Fig. 4e**) showed that the hetrosynaptic LTD decreased with the increase in distance from the stimulus site while homosynaptic LTP was induced. This Mexican hat-like heterosynaptic plasticity was in good agreement with the experimental results (bottom, **Fig. 4f**). For heterosynaptic sites (sites 1–6), the amplitudes of unconstrained energy were always lower than that of the maximum potential energy. Therefore, the maximum potential energy did not have a constraining effect, resulting in the overlap of constrained and unconstrained potential energy (**top**; **Figs. 4b,d,f**). The homosynaptic LTP increase unlimitedly if it was not constrained by energy. This condition was because the unconstrained energy with amplitude greater than the maximum potential energy was not adjusted, and the difference between the baseline and suprathreshold potential energy was extremely large (**middle**; **Figs. 4b,d,f**). The energy constraint made the postsynaptic potential energy equal to the maximum potential energy, thereby reducing the difference between the baseline and suprathreshold potential energy and controlling the homosynaptic LTP in a biologically reasonable range (**bottom**; **Figs. 4b,d,f**). Although the sites of hetero LTD were more than that of homosynaptic LTP, the amplitude of homosynaptic LTP was larger than that of heterosynaptic LTD by comparing homosynaptic LTP and heterosynaptic LTD. As shown in the top panel of **Figures 4b, d**, and **f**, the difference in the maximum potential energy was due to the different signs of the corresponding potential energy, and their energy supply was actually the same.

## Discussion

We presented a computational model of synaptic plasticity completely determined by energy and established a simple quantitative relationship between synaptic plasticity and postsynaptic potential energy. The synaptic weight is directly proportional to the difference between the baseline and suprathreshold potential energy and is constrained by the maximum energy supply. Considering that the dynamic characteristics of energy supply are unclear, we proposed a simple dynamic equation of energy supply and provided the upper limit of the amplitude of postsynaptic potential energy. The constraint of energy supply improves the performance of synaptic plasticity and avoids setting the hard boundary of synaptic weights. In the classical frequency-dependent pairing protocol, six parameters of the model were determined by trial and error. With such a set of parameters, our model reproduced several experimental results of homosynaptic plasticity and the Mexican hat-like heterosynaptic LTD, showing that our model can unify the homo and heterosynaptic plasticity.

### Quantitative relationship between synaptic plasticity and metabolic energy

The absolute value of potential energy can be regarded as the metabolic energy consumed because it restores postsynaptic potential energy to resting state. Our model shows that the synaptic weight is directly proportional to the difference between the baseline and suprathreshold potential energy. Can metabolic energy replace potential energy to express this linear relationship? If the potential energy, baseline, and suprathreshold potential energy are all negative, the absolute value of the baseline potential energy is called the baseline metabolic energy. The absolute value of the suprathreshold potential energy is called the suprathreshold metabolic energy. The weight of the synapse is directly proportional to the difference between the suprathreshold metabolic energy and the baseline metabolic energy. The synaptic weight is linearly related to the metabolic energy if the baseline or suprathreshold potential energy is a constant that does not change with time. Li and van Rossum’s (2020) hypothesis that the metabolic energy is directly proportional to the change in synaptic weight is correct. The linear relationship between synaptic weights and metabolic energy is only valid in some cases.

### Interaction mechanism between Hebbian and homeostatic synaptic plasticity

At present, the biological significance of Hebbian synaptic plasticity (positive feedback) and homeostatic synaptic plasticity (negative feedback) remains controversial. Specifically, how these opposing forms of plasticity that share common downstream mechanisms work in the same networks, neurons, and synapses remain unclear (Turrigiano et al., 1998; Feldman, 2002; Turrigiano and Nelson, 2004; Swanwick et al., 2006; Rannals and Kapur, 2011). In recent years, these conditions have been discussed extensively by leading experts in the field (Vitureira and Goda, 2013; Fox and Stryker, 2017; Keck et al., 2017; Yee et al., 2017). One view is that homeostatic plasticity operates on a long time scale and does not interfere with synaptic changes induced by Hebbian plasticity (Turrigiano, 2012; Tononi and Cirelli, 2014; Hengen et al., 2016). Another view is that Hebbian and homeostatic synaptic mechanisms may be parallel; thus, they can interfere with each other in the same synaptic subset (Keck et al., 2011; Frank, 2014; Li et al., 2014; Desai et al., 2002; Kim and Tsien, 2008; Vlachos et al., 2013). On the basis of observing the existence of fast and input-specific homeostatic mechanisms, a signal pathway-based model was proposed to adjust the balance between Hebbian and homeostatic synaptic plasticity. The model was used to explain the interaction mechanism between Hebbian and homeostatic plasticity on the same time scale (Galanis and Vlachos, 2020).

Our model can integrate the two different viewpoints and give a unified explanation. The homeostatic synaptic plasticity at different time scales coexists. First, we believe that the homeostatic plasticity operating on a long time scale is caused by heterosynaptic plasticity. Our simulation showed that the amplitude of heterosynaptic LTD is extremely small, especially under high-frequency tetanic stimulation (**bottom**; **Figs. 4b,d,f**). This condition indicates that the changes in heterosynaptic plasticity caused by normal neural activities are extremely small or difficult to confirm under the Hebbian time scale. Experiencing a longer neural activity than the Hebbian time scale is necessary before these changes can be evident. This heterosynaptic plasticity accumulated over a long period of time becomes homeostatic plasticity on a long time scale. Second, homeostatic synaptic plasticity on the same time scale as Hebbian synaptic plasticity is caused by the constraint of energy supply. The synaptic strength does not increase continuously under high-frequency stimulation nor does it decrease unlimitedly under low-frequency stimulation due to the constrained energy (bottom; **Figs. 2,3b**). On the contrary, the amplitude of synaptic enhancement or inhibition is reduced to match the energy supply because the energy supply gradually decreases after the duration of neural activity is greater than its time constant (**Equation 2**). The final result is a homeostatic synaptic plasticity parallel to the Hebbian time scale. We propose a unified mechanism for the interaction between Hebbian and homeostatic synaptic plasticity based on the above analysis. The homeostatic homo and heterosynaptic plasticity coexist with homo and heterosynaptic plasticity. The time scale of homeostatic homosynaptic plasticity is the same as that of homosynaptic plasticity (i.e., Hebbian synaptic plasticity), which is rapid and input specific and is caused by the limitation of energy supply. The homeostatic heterosynaptic plasticity has a long time scale, which is caused by the long-term accumulation of heterosynaptic plasticity. Although we do not fully understand the molecular mechanism of heterosynaptic plasticity and energy supply and the actual energy supply dynamics, we believe that the analysis of this mechanism from the cellular level is still valuable.

### Limitations of our approach

First, our synaptic plasticity model can reproduce a series of classical synaptic plasticity experiments by using a detailed biophysical model of a single pyramidal neuron. Although the feasibility of the model can be basically confirmed, further examining the consistency between the model and the experimental results under many stimulation protocols is necessary. Second, in our model parameters, the threshold potential *V_m_* of −60 mV corresponds to the leakage potential *E_L_* (**Methods**) in the biophysical model of neurons. The baseline coefficient *A_r_* of 0.2 indicates that the baseline potential energy is one-fifth of the subthreshold potential energy. Whether the two parameters are universal for different neuron models and what their biophysical significance is remain unclear. Third, the proposed dynamic equation of energy supply (**Eq. 2**) is not supported by experimental data. Can the equation be used as a theoretical prediction to guide future experiments? Can different dynamic equations supported by experiments achieve the same effect of energy constraint in our model? These questions are worthy of further exploration. Finally, the interaction between Hebbian and homeostatic plasticity for large-scale neural networks has an important influence on the learning and memory ability of neural networks. We did not study the validity and scalability of the model in the neural network environment, especially in large-scale neural networks, which is an important direction of our future work.

## Methods

### Model of neuron and synapse

All simulations in this study were conducted on Brian 2 neuron simulator in Python (Goodman and Brette, 2009). We used the model of the biophysical neurons and synapses developed by Bono and Clopath (2017) on Brian 2. Given that L5 and L2/3 pyramidal neuron models are found to have similar results, we only conducted simulation studies on synaptic plasticity in L5 pyramidal neurons (Bono and Clopath, 2017). L5 pyramidal neurons are composed of a spherical soma, an axon, and many dendrite branches, with a total of 1181 compartments. Leakage potential *E_L_* and resting potential of each compartment are −60 and −69 mV, respectively. In accordance with the neuron and synaptic model of Bono and Clopath (2017), we can obtain the membrane potential *V_m_* and membrane current density *I_m_* of each compartment under any stimulation protocol. The unit of membrane potential is mV, and the unit of membrane current density is ampere/meter^2^, abbreviated as A/m^2^. We chose the unit of membrane current density as pA/μm^2^ equivalent to A/m^2^ because the geometric size of neurons in Brian 2 is usually expressed in μm.

### Synaptic plasticity

The variables in Brian 2 simulator are usually expressed and calculated in the form of differential equations. Thus, our synaptic plasticity model needs to be realized in the form of differential equations.

The energy supply (**Eq. 2**) is calculated by using two differential equations. The differential equation of exponential decay factor *S_damp_* = *e^−t/τ^* is expressed as *dS_damp_/dt* = −*S_damp_/τ* with an initial value of one. The part of energy supply that increases linearly with time, *S_lin_* = *R t* is expressed as *dS_lin_/dt* = *R*, and the initial value is zero. Therefore, the energy supply of *t* is *S* = *S_damp_ S_lin_* + *S_0_*.

Postsynaptic potential energy *P* is expressed as the integration of postsynaptic unit membrane power to time and is constrained by energy supply. The differential expression is as follows:

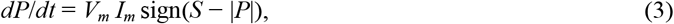

where sign (·) is a symbolic function. The value of the function is −1 when the parameter is negative, and is 1 when the parameter is positive. The value of this function is 0 when it is equal to 0. The unit of *P* is fJ/μm^2^, that is, 10^−15^ J/μm^2^, and the initial value is 0. The unit of *S* and *S_0_* is the same as that of *P*, and the unit of *R* is fJ/(μm^2^ s).

In accordance with the definition and considering the limitation of energy supply, the differential forms of baseline potential energy and suprathreshold potential energy are expressed in *V_m_* and *I_m_* as follows:

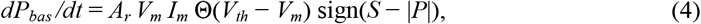

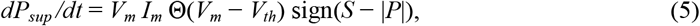

where Θ(·) is the Heaviside step function. The function value is zero when the parameter is negative, otherwise it is one. The initial values of baseline potential energy *P_bas_* and suprathreshold potential energy *P_sup_* are zero.

After substituting **Equations 4** and **5** into **Equation 1**, the differential form of synaptic weights can be expressed completely as follows:

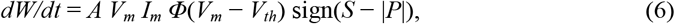

where *Φ*(*V_m_* – *V_th_*) denotes that if *V_m_* < *V_th_*, then *Φ*(*V_m_* – *V_th_*) = *A_r_*, otherwise *Φ*(*V_m_* – *V_th_*) = −1.

## Author Contributions

H.W.C designed the study, designed and implemented the models and wrote the manuscript. L.J.X analyzed the model and participated in discussions. Y.J.W and H.Z. discussed the results and commented on the manuscript.

